# Distinct functions of TIR1 and AFB1 receptors in auxin signalling

**DOI:** 10.1101/2023.01.05.522749

**Authors:** Huihuang Chen, Lanxin Li, Minxia Zou, Linlin Qi, Jiří Friml

**Affiliations:** Institute of Science and Technology Austria (ISTA), 3400 Klosterneuburg, Austria; Beijing Key Laboratory of Development and Quality Control of Ornamental Crops, Department of Ornamental Horticulture, College of Horticulture, China Agricultural University, 100193 Beijing, China

## Abstract

Auxin is the major plant hormone regulating growth and development (Friml, 2022). Forward genetic approaches in the model plant *Arabidopsis thaliana* have identified major components of auxin signalling and established the canonical mechanism mediating transcriptional and thus developmental reprogramming. In this textbook view, TRANSPORT INHIBITOR RESPONSE 1 (TIR1)/AUXIN-SIGNALING F-BOX (AFBs) are auxin receptors, which act as F-box subunits determining the substrate specificity of the Skp1-Cullin1-F box protein (SCF) type E3 ubiquitin ligase complex. Auxin acts as a “molecular glue” increasing the affinity between TIR1/AFBs and the Aux/IAA repressors. Subsequently, Aux/IAAs are ubiquitinated and degraded, thus releasing auxin transcription factors from their repression making them free to mediate transcription of auxin response genes (Yu *et al*., 2022). Nonetheless, accumulating evidence suggests existence of rapid, non-transcriptional responses downstream of TIR1/AFBs such as auxin-induced cytosolic calcium (Ca^2+^) transients, plasma membrane depolarization and apoplast alkalinisation, all converging on the process of root growth inhibition and root gravitropism (Li *et al*., 2022). Particularly, these rapid responses are mostly contributed by predominantly cytosolic AFB1, while the long-term growth responses are mediated by mainly nuclear TIR1 and AFB2-AFB5 (Li *et al*., 2021; Prigge *et al*., 2020; Serre *et al*., 2021). How AFB1 conducts auxin-triggered rapid responses and how it is different from TIR1 and AFB2-AFB5 remains elusive. Here, we compare the roles of TIR1 and AFB1 in transcriptional and rapid responses by modulating their subcellular localization in Arabidopsis and by testing their ability to mediate transcriptional responses when part of the minimal auxin circuit reconstituted in yeast.

**Short summary:** Auxin receptors TIR1 and AFB1 have distinct functions in mediating transcriptional and non-transcriptional responses, respectively. Manipulation of their subcellular localizations revealed that these functional differences cannot be attributed to nuclear versus cytosolic enrichment of TIR1 and AFB1 but to different, specific properties of these proteins, not least to their ability to associate with ubiquitin ligase components.

---

One prominent difference between TIR1 and AFB1 is their subcellular localization. TIR1 primarily localizes to the nucleus, while AFB1 to the cytoplasm (Figure 1A, Supplemental Figure 1) (Prigge *et al*., 2020). To test whether their specific localization is a necessary prerequisite for their function in either transcriptional or rapid responses, we fused *Venus* report gene combined with nuclear exporting signal (NES) or nuclear localization signal (NLS) at the C-terminal of *TIR1* (*TIR1-NES-Venus*) and *AFB1* (*AFB1-NLS-Venus*), respectively. We showed that the majority of TIR1-NES-Venus is shifted to cytosol, while AFB1-NLS-Venus mostly concentrates in the nucleus (Figure 1B, Supplemental Figure 1).

**Figure 1.**
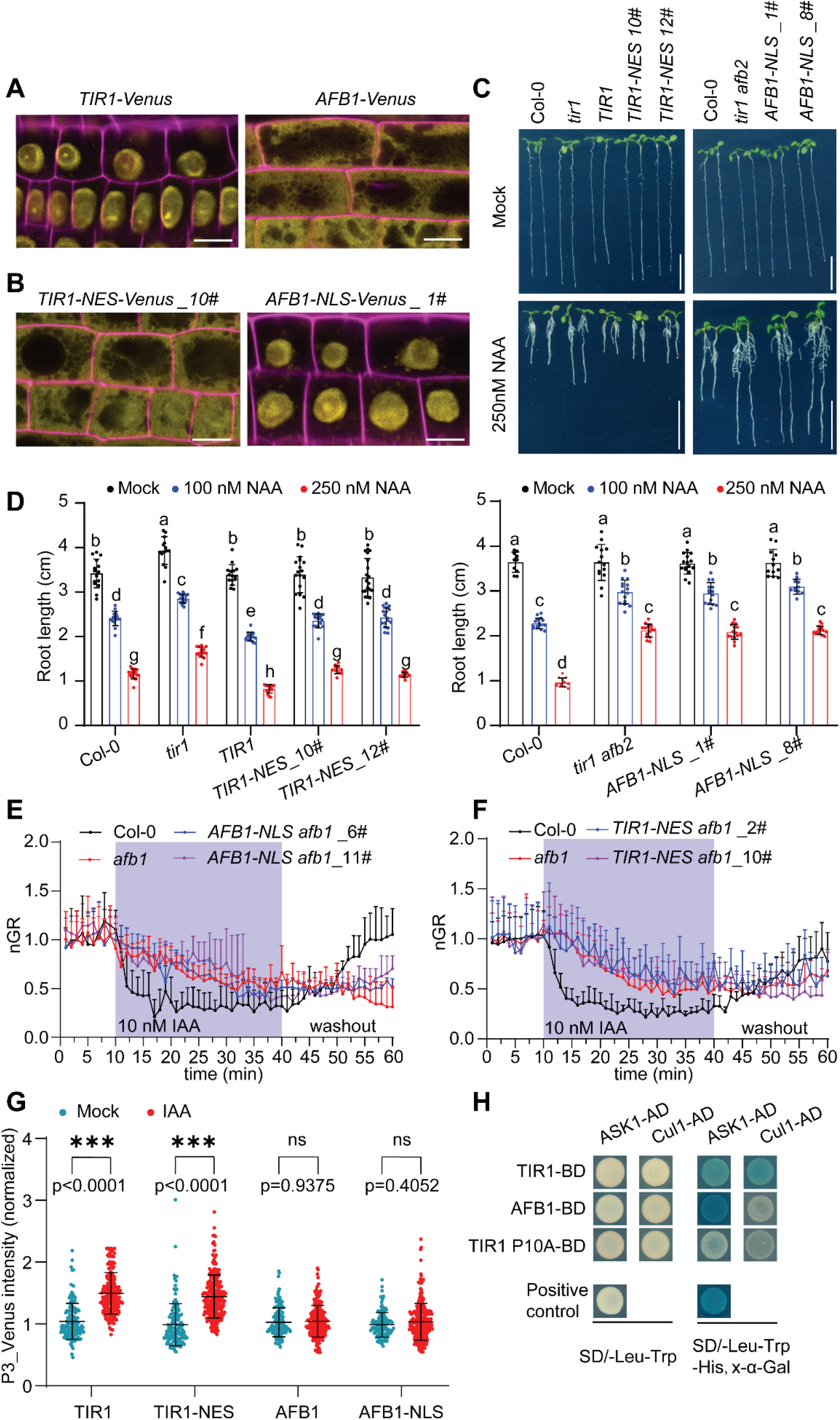
Distinct functions of TIR1 and AFB1 receptors in auxin signalling. (A, B) Confocal images of *Arabidopsis* root epidermis cells expressing *TIR1-Venus, AFB1-Venus, TIR1-NES-Venus* and *AFB1-NLS-Venus* as indicated. Venus signal is shown in yellow in all panels. Cell wall stained with propidium iodide is shown in purple. Scale bars =10 μm. (C, D) Evaluation of the roles of TIR1-NES-Venus and AFB1-NLS-Venus in auxin-mediated long-term root growth inhibition. The indicated *TIR1*-related constructs were transformed into *tir1*, and the *AFB1*-related constructs were transformed into *tir1 afb2*. Primary root length in the 6-day-old seedlings of different genotypes grown on mock or 250nM NAA plate. Scale bars = 1 cm, n ≥ 10 (D). Lowercase letters indicate significant difference, two-way ANOVA test, *p*≤0.01. (E, F) Evaluation of the roles of TIR1-NES-Venus and AFB1-NLS-Venus in in auxin-mediated rapid root growth inhibition using Microfluidic vRootchip. The constructs *TIR1-NES-Venus* and *AFB1-NLS-Venus* were transformed into *afb1-3* mutant. Root growth rate of the indicated genotypes was normalized to the respective average root growth rate within 10 minutes before IAA application. The imaging interval is 1 minute. (G) Differential effects of TIR1, TIR1-NES, AFB1, AFB1-NLS in auxin mediated transcriptional responses in the minimal auxin signaling pathway reconstructed in *Saccharomyces cerevisiae*. Fluorescence intensity was quantified with the images captured after the yeast cells were treated with 10 μM IAA for 6 hours. Error bar = ± SD, *t* test, \*\*\**P*□≤□0.001, NS, not significant (*P*>0.05). (H) Interactions of different SCF components in yeast two hybrid assay. The yeast transformants were plated on SD/-Leu-Trp-His drop out medium with 4 mg/ml x-alpha-Gal (x-α-Gal) and were cultured for 3 days to assess the protein-protein interactions. TIR1 P10A acts as a negative control in here. Alpha-galactosidase activity manifested as blue color indicates the interaction.

To characterize the importance of nuclear versus cytosolic localization of TIR1 and AFB1 in auxin-mediated transcription, we tested if they can rescue mutant phenotype in a sustained root growth inhibition after auxin treatment for 6 days. We introduced the cytosolic-localized *TIR1-NES-Venus* into *tir1* mutant background, and found that all *TIR1* constructs were able to restore completely auxin sensitivity of root growth (Figure 1C, D and Supplemental Figure 1, 2). This can be explained by either: (i) the residual TIR1 present in the nucleus is sufficient to conduct full transcriptional activity, or (ii) cytosolic TIR1 may still degrade Aux/IAAs, releasing the ARFs from their inhibition. Besides, we introduced the nuclear-localized *AFB1-NLS-Venus* into *tir1 afb2* mutants but did not observe any rescue of the auxin-insensitive phenotype in root growth inhibition, root gravitropism, lateral root formation, and root hair elongation (Figure 1C, D and Supplemental Figure 1, 2). This implies that AFB1, even when localized to nucleus, cannot functionally replace TIR1 for its transcriptional regulation and related development.

The predominantly cytosolic AFB1 seems to be the major receptor for the rapid auxin effects (Prigge *et al*., 2020). Therefore, we introduced our mistargeted TIR1 and AFB1 versions into the *afb1* mutant background (Figure 1A, B and Supplemental Figure 1) and test their effect in auxin-induced rapid root growth inhibition in microfluidic vRootchip system. The AFB1 when targeted to nucleus could no longer mediate rapid auxin effect on root growth (Figure 1E). On the other hand, TIR1 despite being present in the cytosol could not rescue *afb1* mutant (Figure 1F). This reveals that the cytosolic AFB1 is necessary for its function, but the cytosolic TIR1 cannot replace or supplement the AFB1 function.

The observations that nuclear AFB1 cannot functionally replace TIR1 and cytosolic TIR1 cannot functionally replace AFB1 show that TIR1 and AFB1 have distinct functional properties unrelated to their subcellular localization. To confirm this, we made use of the minimal auxin signalling pathway reconstructed in yeast (Pierre-Jerome *et al*., 2014). In this system, only TIR1 but not AFB1, regardless of their subcellular localization, was able to mediate auxin effect on transcription as monitored by the fluorescence intensity of P3_Venus transcriptional auxin reporter (Figure 1G).

To gain insights into reasons why AFB1 cannot mediate transcriptional signalling, we tested its ability to form SCF complex using Yeast-Two-Hybrid approach. Only TIR1 but not AFB1 was able to interact with CUL1 (Cullin1), the key component of ubiquitin ligase complex (Figure 1H). This is consistent with the available Co-IP/MS data where all SCF components were detected to interact with TIR1, however, for AFB1, no or only extremely weak interaction with CUL1 was detected (Supplemental Figure 3) (Li *et al*., 2021; Yu *et al*., 2015). The reason why AFB1 does not interact with CUL1 might be the natural mutation of glutamic 8 site in AFB1 (Yu *et al*., 2015). The absence of interaction with the SCF components will prevent AFB1 to conduct E3 ubiquitin ligase activity; thus failing to mediate Aux/IAAs degradation (Yu *et al*., 2015) and transcriptional regulation (Figure 1G). This also explains why AFB1, even when artificially targeted to nucleus, still cannot replace the TIR1 function.

Our observations also imply that CUL1 is not essential for rapid auxin responses. This requires further clarification on the role of SCF components as well as Aux/IAAs ubiquitination and degradation in rapid auxin responses. Recent study revealed the novel function of TIR1/AFBs in producing cAMP, a prominent second messenger in animals. Though this activity in TIR1 specifically does not seem to be important for rapid auxin responses (Qi *et al*., 2022), it is still possible that AFB1-mediated cAMP production in cytosol is. However, whether and how the AC activity of AFB1 would contribute to rapid responses remains unknown.

In summary, we demonstrated that TIR1 and AFB1 have distinct functions with the predominantly nuclear TIR1 mediating slow responses and cytosolic AFB1 conducting the rapid responses. This functional divergence is not, however, simply due to differential subcellular localization of these auxin receptors. The function of TIR1 in mediating slow/transcriptional response seems to be independent of its predominant localization. In contrast, the function of AFB1 in rapid responses necessitates both its localization in cytosol and the specific AFB1 protein properties themselves. Furthermore, the cytosolic AFB1 mediates rapid auxin responses without forming SCF machinery, leaving the mechanism of AFB1-mediated rapid responses an exciting topic for future investigations.

## SUPPLEMENTAL INFORMATION

### Supplemental Method

**Supplemental Figure 1**. Subcellular localization of *AFB1-Venus, TIR1-NES-Venus*, and *AFB1-NLS-Venus* in different transgenic lines.

**Supplemental Figure 2**. Evaluation of the roles of *TIR1-NES-Venus* and *AFB1-NLS-Venus* in auxin-mediated long-term responses.

**Supplemental Figure 3**. IP-MS/MS results performed with the *pTIR1::TIR1-Venus* and *pAFB1::AFB1-Venus* lines.

## AUTHOR CONTRIBUTIONS

H.C., L.L. and J.F. designed the studies; H.C. and M.Z. performed the experiment. H.C., L.Q. and J.F. wrote the manuscript.

## ACKNOWLEDGMENTS

We thank all the authors for sharing the published materials. This research was supported by the Lab Support Facility and the Imaging and Optics Facility of ISTA. We thank Lukáš Fiedler (ISTA) for critical reading of the manuscript.

## The authors declare no competing interests

## Supplemental Method

### Plant materials and growth conditions

All *Arabidopsis* mutants and transgenic lines used in this study are in the Columbia-0 (Col-0) background. The *tir1-1* (Ruegger et al., 1998), *afb1-3* (Savaldi-Goldstein et al., 2008), *tir1-1 afb2-3* (Prigge et al., 2020), *pTIR1::TIR1-Venus* in *tir1-1* (Wang et al., 2016) and *pAFB1::AFB-Venus* in *afb1-3* (Rast-Somssich et al., 2017) mutants were donated by the original authors. The promoters of *TIR1* (Wang *et al*., 2016) and *AFB1* (Rast-Somssich et al., 2017) were amplified from genomic DNA and cloned into pDONR P4-P1r Gateway entry vector. The genomic or coding domain sequences of TIR1 and coding domain sequences of AFB1 without stop codon were cloned into pENTR/D-TOPO Gateway entry Vector to obtain the pENTR-gTIR1, pENTR-TIR1 and pENTR-AFB1. The NES-GS-linker (ctgcagctgcctcccctggagcgcctgaccctggacggaggcggtggaagcggcggaggt) and NLS-GS-linker (atgcccaagaagaagagaaaggtaggaggcggtggaagcggcggaggttcc) were directly introduced at the 5’ end of Venus by primers and cloned into the pDONR-P2r-P3 vector. All those plasmids were confirmed by sequencing and then recombined into the destination vector pB7m34GW,0 by LR reaction. The final expression plasmids were introduced into *Agrobacterium tumefaciens* GV3101 strain by electroporation. *Arabidopsis* plants were transformed by floral dip. Two independent homozygous lines for each transgenic plant were used for further experiments. Seed surface-sterilization and growth conditions were the same as described before (Qi et al., 2022).

### Phenotypic analysis

For root length measurement: the seeds from the same batch were sown directly on the ½ Murashige and Skoog (MS) medium with 100 nM or 250 nM NAA. Medium with equally diluted ethanol was used as the Mock group. The plates containing 6-day-old seedlings were imaged by a scanner. The primary root length was measured manually by using the segmented Line plug-in of Image J.

For gravitropism analysis: 5-day-old seedlings were transferred to a new plate containing ½ MS medium, and the plate was rotated 90 degrees for gravity stimulation and was placed on a vertical scanner. Images were automatically taken using Autolt program every 30 min, and the root growth deviation angle was measured using Image J.

For lateral root formation: 5-day-old seedlings were transferred to ½ MS medium containing mock or 100 nM IAA, and grown for another 5 days. The number of lateral roots were counted from the images obtained.

For root hair length measurement: 5-day-old seedlings were transferred to ½ MS medium containing mock or 10 nM IAA, and grown for 24 h. After taking images with a stereo microscope (Olympus SZX16), root hair length was measured using Image J.

### Microfluidic vRootchip

The upgraded Microfluidic vRootchip system was used to analyze the auxin-mediated rapid root growth inhibition (Li et al., 2021). The 4-day-old seedlings were transferred into the vRootchip channel and grown in ¼ MS medium without sucrose for more than 12 hours. The vRootchip was then mounted on the vertical confocal microscope system and the medium was replaced with ¼ MS medium containing 0.1% sucrose (Von Wangenheim et al., 2017). The imaging started after the plants were acclimated for 3 hours. Zeiss LSM 800 confocal microscope with transmitted light detector was used for imaging. The objective lens was 20×/0.8 NA air objective, and the Zoom in was 1.0 ×. The time interval for imaging was 1 minute. After pre-imaging for 10 minutes, medium containing 10 nM IAA was applied, followed by a washout with basic medium after 30 minutes. The software package used for data analysis is the same as described before (Li et al., 2021).

### Minimal auxin signalling pathway constructed in yeast

The vectors, plasmids, and yeast strains for this assay were all provided by Prof. Jennifer, including pGP5G-ccdb empty vector, pGP4GY-TPLN100-IAA3 plasmid, pGP8A-ARF19 plasmid, MATa strain containing the P3_2x-UbiVenus reporter integrated at URA3, and MATa strain containing the HIS3:pADH1-ARF19 URA3:P3_2x-UbiVenus (Pierre-Jerome et al., 2014). pENTR-TIR1, pENTR-TIR1-NES, pENTR-AFB1 and pENTR-AFB1-NLS were recombined with the destination vector pGP5G-ccdb to generate pGP5G-TIR1, pGP5G-TIR1-NES, pGP5G-AFB1 and pGP5G-AFB1-NLS plasmids by LR reaction. Yeast transformation was performed by the lithium acetate transfer method (Sherman, 2002). Transformants survived from SD/-Leu-Trp-Ura-His drop out (QDO) solid medium were transferred to QDO liquid medium to grow overnight. Then, the yeast cells were diluted 10 times in QDO liquid medium and treated with 10 μM IAA for 6 h. Images were taken by an inverted Zeiss LSM800 confocal microscope with 20×/0.8 NA air objective. The Venus fluorescence intensity was measured using Image J, with more than 100 yeast cells for each treatment.

### Yeast two hybrid

Phusion Site-Directed Mutagenesis Kit (Thermo Fisher, F541) was used to generate the P10A mutation in TIR1, using pENTR-TIR1 as the template. pENTR-TIR1, pENTR-TIR1 P10A and pENTR-AFB1 were introduced into pGBKT7-GW vector (Clontech). The coding domain sequence of ASK1 and Cullin1 were cloned into the pGADT7 vector (Clontech). Different pGBKT7 recombinant plasmids and pGADT7 recombinant plasmids were co-transformed into Y2H Gold yeast strain (Clontech) by lithium acetate transformation (Sherman, 2002). SD/-Leu-Trp drop-out (DDO) plate is used to confirm the presence of the transgene. The transformants with the size of 2-3 mm were suspended in DDO liquid medium and incubated at 30 °C until OD_600_ was around 0.8. 3 μl of suspended culture was dropped on the SD/-Leu-Trp-His plate containing 4mg/ml X-alpha-Gal. Alpha -galactosidase activity is used to evaluate the interaction after incubation at 30 °C for 3 days.

---

**Supplemental Figure 1.**
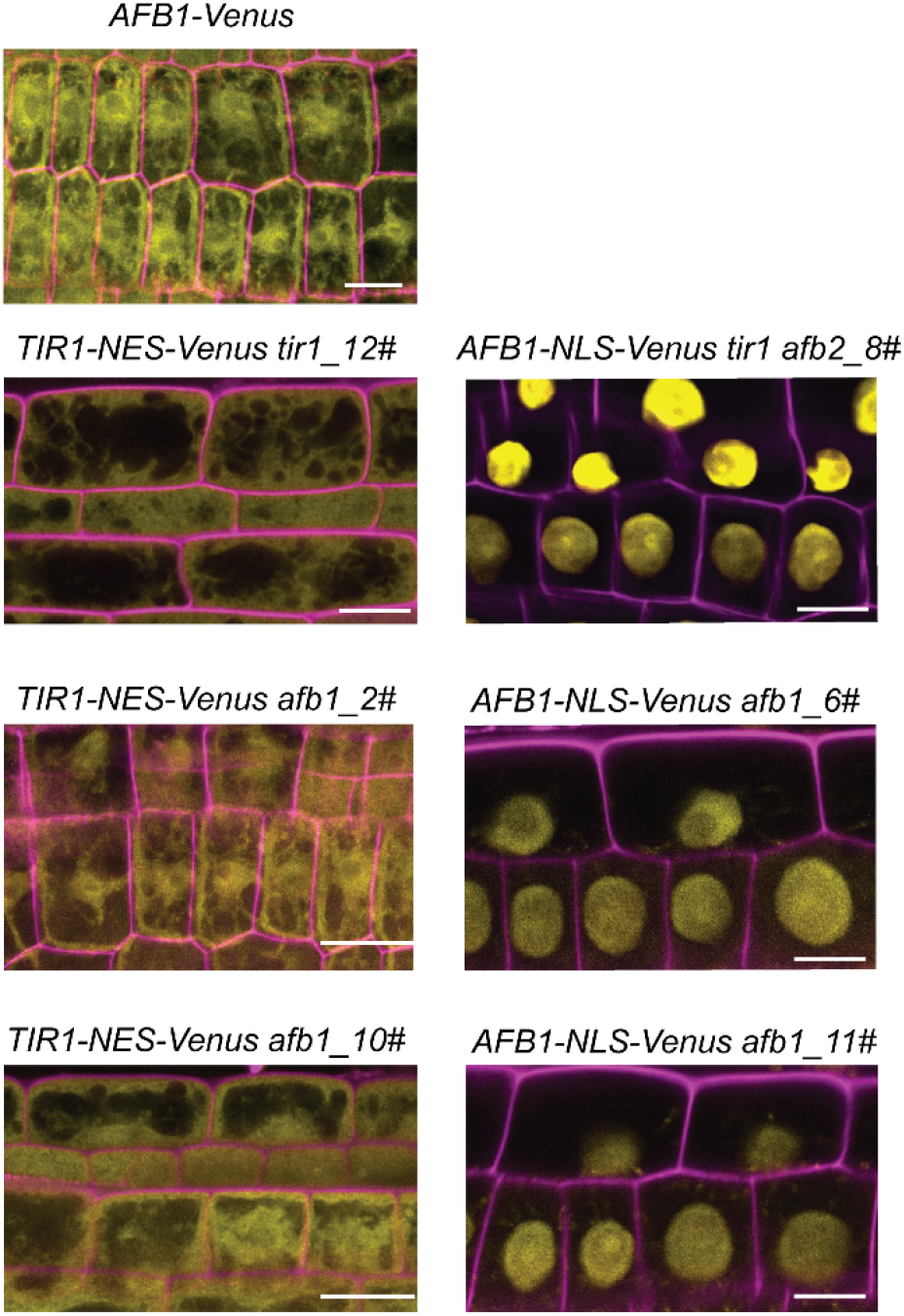
Subcellular localization of *AFB1-Venus, TIR1-NES-Venus* and *AFB1-NLS-Venus* in different transgenic lines. Confocal images of the root epidermis cells of the 4-day-old *Arabidopsis* seedlings with the indicated genotypes: *AFB1-Venus, TIR1-NES-Venus* in *tir1-1* or *afb1-3, AFB1-NLS-Venus* in *tir1-1 afb2-3* or *afb1-3*. Venus signal was shown in yellow in all panels. Cell wall was stained with propidium iodide and shown in purple. The lines shown here were also used in Figure1C, E and F. The *AFB1-Venus* shows weak nuclear signal in certain Z-plane. Scale bars = 10 μm.

**Supplemental Figure 2.**
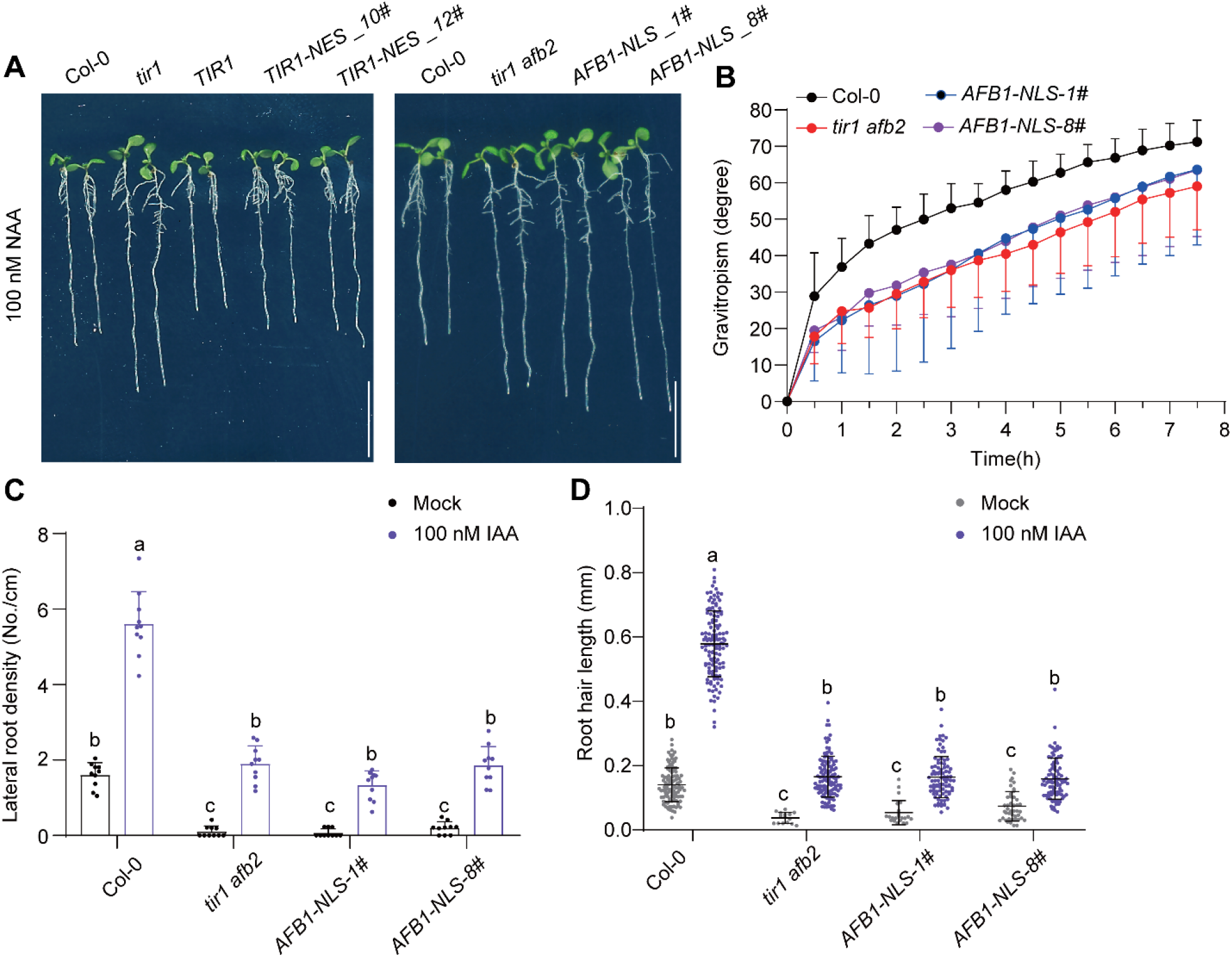
Evaluation of the roles of *TIR1-NES-Venus* and *AFB1-NLS-Venus* in auxin-mediated long-term responses. **(A)** Shown were 6-day-old seedlings of the indicated genotypes grown on the ½ MS medium containing 100 nM NAA. Scale bars = 1 cm. (B-D) AFB1-NLS cannot exert TIR1 function in root gravitropism, later root formation, and root lair elongation. (B)Five-day-old seedlings were transferred to new plates and were then rotated 90°. Images were captured every 1 h, and the root bending angle was measured to monitor the gravitropic response. Data are mean□+/-□SD of 10 seedlings. (C) Five-day-old seedlings were transferred to mock or 100 nM IAA-containing medium and were cultured for another 5 days. Lateral root density was calculated from 10 roots. Error bars = + SD. Different lowercase letters indicate significant difference, *p*≤□0.001. (D) Five-day-old seedlings were transferred to mock or 10 nM IAA medium. The root hair length was measured after growth for 24 h. Error bars = ±□SD, Different lowercase letters indicate significant difference, *p*≤□0.001.

**Supplemental Figure 3.**
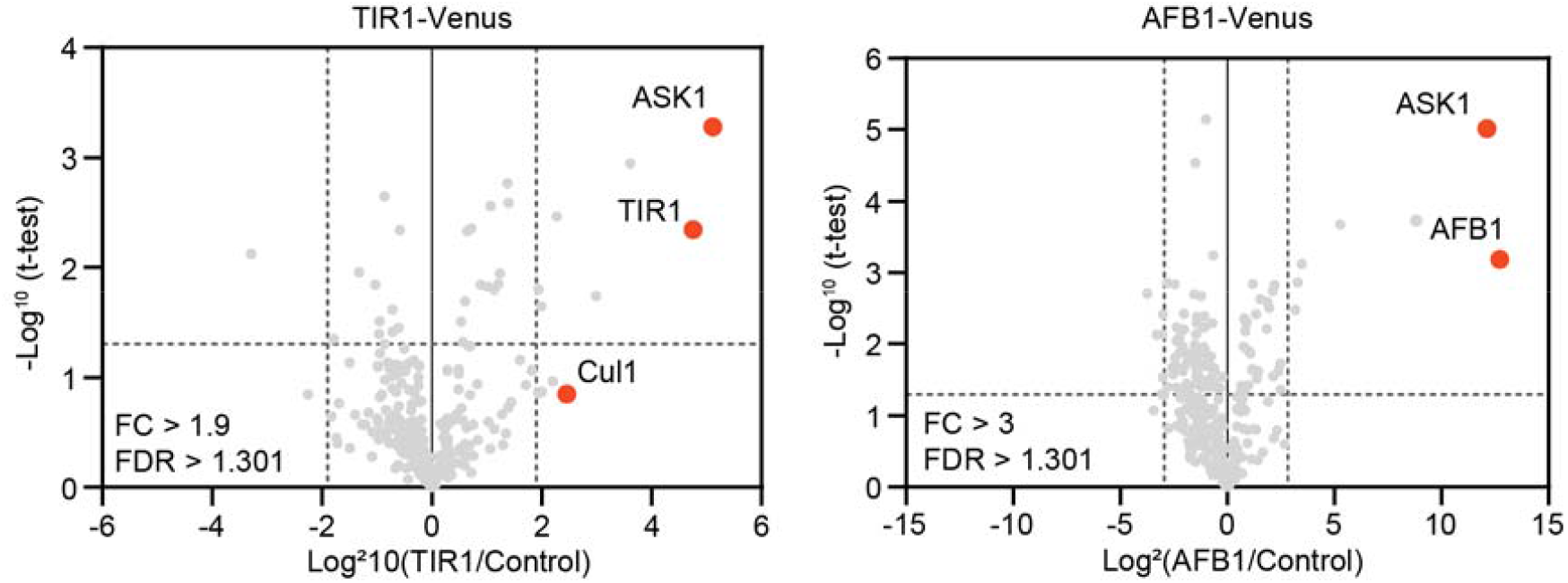
IP-MS/MS results performed with the *pTIR1::TIR1-VENUS* and *pAFB1::AFB1-VENUS* lines. Red dots indicate the subunits of SCF complex. Note that CUL1 was not detected in the immune precipitates of AFB1-Venus. The original data was available in Li et al, 2022, Supplementary Table 2.

## References

Friml, J. (2022). Fourteen Stations of Auxin. Cold Spring Harb Perspect Biol 14. 10.1101/cshperspect.a039859.

Li, L., Gallei, M., and Friml, J. (2022). Bending to auxin: fast acid growth for tropisms. Trends in Plant Science 27, 440–449. 10.1016/j.tplants.2021.11.006.

Li, L., Verstraeten, I., Roosjen, M., Takahashi, K., Rodriguez, L., Merrin, J., Chen, J., Shabala, L., Smet, W., Ren, H., et al. (2021). Cell surface and intracellular auxin signalling for H(+) fluxes in root growth. Nature 599, 273–277. 10.1038/s41586-021-04037-6.

Pierre-Jerome, E., Jang, S.S., Havens, K.A., Nemhauser, J.L., and Klavins, E. (2014). Recapitulation of the forward nuclear auxin response pathway in yeast. Proceedings of the National Academy of Sciences 111, 9407–9412. 10.1073/pnas.1324147111.

Prigge, M.J., Platre, M., Kadakia, N., Zhang, Y., Greenham, K., Szutu, W., Pandey, B.K., Bhosale, R.A., Bennett, M.J., Busch, W., and Estelle, M. (2020). Genetic analysis of the Arabidopsis TIR1/AFB auxin receptors reveals both overlapping and specialized functions. Elife 9. 10.7554/eLife.54740.

Qi, L., Kwiatkowski, M., Chen, H., Hoermayer, L., Sinclair, S., Zou, M., Del Genio, C.I., Kubeš, M.F., Napier, R., Jaworski, K., and Friml, J. (2022). Adenylate cyclase activity of TIR1/AFB auxin receptors in plants. Nature 611, 133–138. 10.1038/s41586-022-05369-7.

Serre, N.B.C., Kralík, D., Yun, P., Slouka, Z., Shabala, S., and Fendrych, M. (2021). AFB1 controls rapid auxin signalling through membrane depolarization in Arabidopsis thaliana root. Nature Plants 7, 1229–1238. 10.1038/s41477-021-00969-z.

Yu, H., Zhang, Y., Moss, B.L., Bargmann, B.O., Wang, R., Prigge, M., Nemhauser, J.L., and Estelle, M. (2015). Untethering the TIR1 auxin receptor from the SCF complex increases its stability and inhibits auxin response. Nat Plants 1. 10.1038/nplants.2014.30.

Yu, Z., Zhang, F., Friml, J., and Ding, Z. (2022). Auxin signaling: Research advances over the past 30 years. Journal of Integrative Plant Biology 64, 371–392. 10.1111/jipb.13225.

## Reference

Rast-Somssich, M.I., Žádníková, P., Schmid, S., Kieffer, M., Kepinski, S., and Simon, R. (2017). The Arabidopsis JAGGED LATERAL ORGANS (JLO) gene sensitizes plants to auxin. Journal of Experimental Botany 68, 2741–2755. 10.1093/jxb/erx131.

Ruegger, M., Dewey, E., Gray, W.M., Hobbie, L., Turner, J., and Estelle, M. (1998). The TIR1 protein of Arabidopsis functions in auxin response and is related to human SKP2 and yeastLGrr1p. Genes & Development 12, 198–207. 10.1101/gad.12.2.198.

Savaldi-Goldstein, S., Baiga, T.J., Pojer, F., Dabi, T., Butterfield, C., Parry, G., Santner, A., Dharmasiri, N., Tao, Y., Estelle, M., et al. (2008). New auxin analogs with growth-promoting effects in intact plants reveal a chemical strategy to improve hormone delivery. Proceedings of the National Academy of Sciences 105, 15190–15195. 10.1073/pnas.0806324105.

Sherman, F. (2002). Getting started with yeast. In Methods in Enzymology, C. Guthrie, and G.R. Fink, eds. (Academic Press), pp. 3–41. 10.1016/S0076-6879(02)50954-X.

Von Wangenheim, D., Hauschild, R., Fendrych, M., Barone, V., Benková, E., and Friml, J. (2017). Live tracking of moving samples in confocal microscopy for vertically grown roots. eLife 6. 10.7554/elife.26792.

Wang, R., Zhang, Y., Kieffer, M., Yu, H., Kepinski, S., and Estelle, M. (2016). HSP90 regulates temperature-dependent seedling growth in Arabidopsis by stabilizing the auxin co-receptor F-box protein TIR1. Nature Communications 7, 10269. 10.1038/ncomms10269.

